# Integrating Ideal Bayesian Searcher and Neural Networks Models for Eye Movement Prediction in a Hybrid Search Task

**DOI:** 10.1101/2024.11.29.626088

**Authors:** Gonzalo Ruarte, Damián Care, Gaston Bujia, Matias J Ison, Juan E. Kamienkowski

**Author notes:** Both authors equally contributed to the present work (Matias J Ison, Juan E. Kamienkowski).

## Abstract

Visual search, where observers search for a specific item, is a crucial aspect of daily human interaction with the visual environment. Hybrid search extends this by requiring observers to search for any item from a given set of objects. While there are models proficient at simulating human eye movement in visual search tasks within natural scenes, none are able to do so in Hybrid search tasks within similar environments. In this work, we present an enhanced version of the neural network Entropy Limit Minimization (nnELM) model, which is based on a Bayesian framework and decision theory. We also present the Hybrid Search Eye Movements (HSEM) Dataset, comprising several thousands of human eye movements during hybrid search tasks in natural scenes. A key challenge in Hybrid search, absent in visual search, is that participants might search for different objects at different time points. To address this, we developed a strategy based on the posterior probability distribution generated after each fixation. By adjusting the model’s peripheral visibility, we made early search stages more efficient, aligning it closer to human behaviour. Additionally, limiting the model’s memory capacity reduced its success in longer searches, mirroring human performance. To validate these improvements, we compared our model against participants from the HSEM dataset and against existing models in a visual search benchmark. Altogether, the new nnELM model not only successfully explains Hybrid search tasks, but also closely replicates human behaviour in natural scenes. This work advances our understanding of complex processes underlying visual and Hybrid search while maintaining model interpretability.

## 1 Introduction

Visual search is a critical cognitive process involved in many different daily tasks. For instance, every morning when preparing breakfast, we have to look for a cup, coffee, a spoon and so on. At the core of this task are the various eye movements, of which the most important ones are fixations and saccades. A fixation occurs when the eye remains focused on a particular location, typically for 200 ms. When this happens, new visual information about the scene is acquired. The rapid movements that occur between fixations, known as saccades, occur too quickly to provide new visual information but are essential to move the eyes towards a different location in the scene. The ordered sequence of fixations, known as a scanpath, is known to depend on several characteristics including the task performed [43], as well as different features from the perceived scene [37, 32].

Real-life searches often involve searching for multiple objects listed in memory. Going back to the breakfast example, we typically don’t perform the search in a strict sequence, nor for exact items, and we look for any cup, spoon, plate, and so on. Thus, we need to look both into the scene and into the list of items held in memory (memory search). When observers search for any of several potential targets, the task is called Hybrid search (HS) [38]. First introduced by Schneider and Shiffrin’s seminal work [31], it incorporates both visual search (VS) and memory search (MS).

While in real-life searches humans are looking at scenes (mostly everyday scenes), typically experiments are done with participants looking at images on a computer screen and an eye-tracking device to record their eye movements. To date, the large majority of hybrid search findings are derived from experiments utilizing images of artificial noise or shapes against blank backgrounds [40], potentially limiting their ecological validity [23]. In visual search, an increasing body of evidence emphasizes the significance of context (i.e. a kitchen scenery) in guiding attention during real-world searches [39].

In recent years, some computational models for Visual Search (VS) in natural scenes have emerged, with some primarily designed to emulate behaviour during specific tasks, such as searching for a class of object (e.g. cups), with limited generalization ability [42]. Other approaches focused on establishing a correlation between these models and the cognitive processes associated with the task [5, 45, 29, 44]. Najemnik and Geisler [21] proposed the Ideal Bayesian Searcher model (IBS), using decision theory where each fixation is modelled as a decision on where to look next. This seminal paper led to related approaches in more recent years. Najemnik and Geisler [22] used Entropy Limit Minimization (ELM) instead of Bayesian integration for the decision-making process. These models accumulated evidence with each fixation to update their posterior probability. Zhou and Yu [45] further extended the ELM model to take the fixation duration into account in the evidence accumulation process. They also proposed a way to limit the amount of fixations that are used to decide where to go next. These models were tested on artificial images which did not depict natural scenes. Bujia et. al. [5] proposed a way to extend the IBS to natural scenes. This involves the use of Deep Neural Networks to generate saliency maps for the model’s prior, and employing a similarity metric between the target and the image regions in a natural scene when estimating the model’s template response. This model was further improved in [36] by using the attention map of [44] as template response and in [6] by using ELM instead of the guidance and integration from IBS. Rashidi et. al. [29] developed a new model of target detectability, which was then tested in an IBS in natural scenes. To date, there remains a lack of consensus regarding the most appropriate metrics, although ongoing research endeavours are addressing this gap [36].

Remarkably, there appears to be a lack of computational models dedicated to hybrid search tasks in natural scenes. In the current study, we extend the model presented in [6] to hybrid search tasks. Briefly, we explore different ways of limiting the amount of fixations used to decide where to go next, and we improve the peripheral visibility of the model. Altogether, we achieved higher performance metrics as well as increased computational efficiency.

One of the major bottlenecks limiting comparisons between computational models and human behaviour is the availability of well-documented open datasets. A secondary objective of the current study is to generate a visual and hybrid search dataset comprising actual scanpaths from participants. Typically, participants are instructed to look for objects across a large number of images depicting natural scenes. In this work, we also present the Hybrid Search Eye Movements (HSEM) dataset, which will be available after publication.

We use a fixed subset of our hybrid search dataset to test the various modifications to the model, and its hybrid search capabilities. Furthermore, the final version is validated on a different subset of the same dataset. Moreover, we gauge its performance against state-of-the-art visual search models in the ViSioNS benchmark. The overall goal of the present work is to advance in the path of bringing these models closer to humans in a wider range of tasks while maintaining interpretability.

## 2 Methods

### 2.1 Hybrid Search Eye Movements (HSEM) Dataset

#### 2.1.1 Participants

The participant pool consisted of two groups: 18 individuals from UBA (Universidad de Buenos Aires) and 28 from UoN (University of Nottingham). Two from the UoN were discarded due to a poor overall performance (less than 55% of accuracy). This gives a total of 44 participants (18 UBA and 26 UoN). The final sample involved 23 male and 21 female participants between 19 and 40 years old (24.5 ± 5.3 years old). All participants were naive to the experiment’s objectives, possessed normal or corrected-to-normal vision, and willingly provided written informed consent. Ethical approval for the study was obtained from the respective Ethics Committees of each university (Protocol 284 from the Instituto de Investigaciones Médicas “Alfredo Lanari” – University of Buenos Aires, and Protocol F1317 from the University of Nottingham).

#### 2.1.2 Stimuli

A total of 210 stimuli (or images) of 1280×1024 pixels were prepared. As shown in Figure 1A, they were constructed by superimposing real-world background images (depicting outdoor scenes –e.g. forest-, and indoor scenes –e.g. shelf-) with 16 individual items (objects, animals or full-body humans). These 16 individual items comprised the visual set of the stimulus. Items in the visual set of one stimulus did not appear in the visual set of other stimuli, with the exception of 53 items that appeared in two separate visual sets, and 5 items that appeared in three different visual sets. This yielded a total of 1569 items across visual sets.

**Figure 1.**
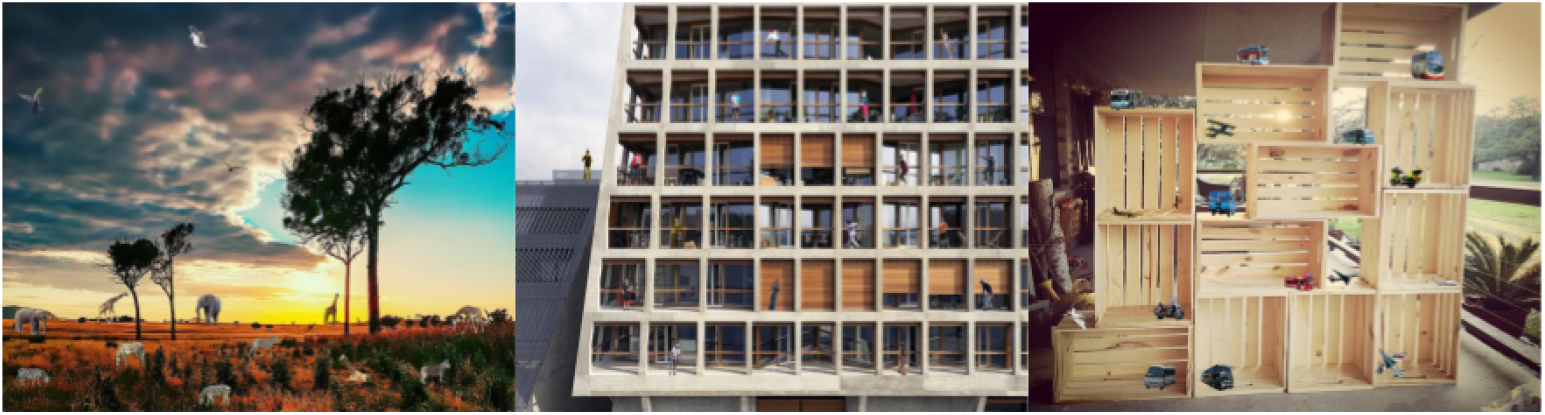
Hybrid Search Eye Movements (HSEM) Dataset example stimuli.

Every item and background image was taken from COCO [20] or ImageNet [10] datasets. The items were resized so that they would fit in the background image, and placed according to scene syntax (i.e. no major violations of support, interposition, position, and size). After the items were placed, a bounding box was defined for each of them which occupied a mean of 0.33% +-0.11% of the background image (Fig 4 B & C). The maximum size a bounding box could have was 80×80 pixels.

For each stimulus, a set of 1, 2, or 4 items were selected for memorization (memory set). In this context, a trial was defined as a stimulus plus its memory set. Within each trial, both the items in the memory set and the ones in the visual set belonged to the same category (objects, animals, or people). Moreover, items in the memory set of a given trial did not appear in the memory set of other trials. There were two exceptions to this rule: two items appeared in the memory sets of two different trials. The visual and memory sets of a given trial could share at most one item: If they did share an item the trial was labelled as a target-present trial–with the shared item being the target–, and the trial was labelled as a target-absent trial otherwise. Targets were unique among target-present trials.

Only the target-present trials were considered for the current analysis. Three of them were eventually discarded from the analysis because their stimuli had a different aspect ratio from the rest. This yielded a total of 102 trials: 35 with a memory set size (MSS) of 1, 34 with a memory set size (MSS) of 2, and 33 with a memory set size (MSS) of 4.

### 2.1.2 Task

Initially, participants were exposed to the memory set for 2 seconds if the MSS was 1, for 3.5 seconds if the MSS was 2, and for 5 seconds if the MSS was 4. To do so, four locations in the screen were previously defined. If the center of the screen was (0,0) then the four locations were: (-300,0), (-150,0), (150,0) and (300,0) in pixels. For each participant and trial, the items in the memory set were randomly placed in one of these locations accompanied by random vertical shifts of 0, ± 50 or ± 100 pixels. Subsequently, participants viewed the stimulus and were allotted 7 seconds to locate the target object (Fig. 4 A). They were required to press one key if they identified the target and a different key if they believed the target was absent. The experiment was divided into seven blocks of 30 trials and different participants were presented with a different order of trials. Every participant had to do each trial only once. The experiments were implemented and executed in Psychopy [26].

### 2.14 Data collection

All the equipment used in the acquisition was the same at both universities except for the use of a ‘qwerty’ keyboard to introduce responses at UBA and a response pad (check) at UoN.

The stimuli were presented in a BenQ XL2420Z monitor at UBA and in a LENOVO Y27Q-20 monitor at UoN with a screen resolution of 1920 x 1080 pixels. Participants were placed at a distance of approximately 65 cm from the monitor. All stimuli were presented using Psychopy software [26] and synchronized with the eye tracker using ioHub^1^.

Eye-movement tracking (ET) data was recorded using an EyeLink 1000 Plus remote system in monocular mode and a sampling rate of 500 Hz. At UoN an EyeLink target sticker was used to ensure movement stabilization while a chinrest was used at UBA for head-stabilization.

There were issues in the collected data of 24 participants so some of their trials are missing. One participant had 12 trials missing but the other 23 had 1 or 2 trials missing at most. This meant a total of 4442 valid trials, 1520 for MSS 1, 1485 for MSS 2 and 1437 for MSS 4.

### 2.1.5 Eye movement pre-analysis

Fixations, saccades and blinks were detected using the built-in EyeLink algorithm. A preprocessing as described in Travi, Ruarte, et al (2022) followed the detection process:

~~~
   **Step 1: Check if the Target is Found by a Fixation
   for each fixation in the scanpath do
   if fixation lands on target bounding box (within 68 pixels) then**
       Mark target as found
       **Go to Step 4
     end if
   end for
   Step 2: Lump Consecutive Close Fixations
   for each fixation in the scanpath do
     for next fixations in the scanpath do
         if distance between current to next fixation *<* 68 pixels then**
           Combine current fixation with next fixation (average positions)
           Remove next fixation from scanpath
        **end if
     end for
   end for
   Step 3: Adjust Fixations Outside Image Boundaries
   for each fixation in the scanpath do
       if fixation coordinates are outside image boundaries then
          Move fixation to nearest point within image
      end if
   end for
   Step 4: Terminate the Scanpath at Target Found or Upper Saccade Limit
   if target is found then**
       Cut scanpath at current fixation
   **else if number of fixations *>* saccade limit then**
       Cut scanpath at saccade limit
   **end if**
   End scanpath processing
~~~

### 2.2 Models

#### 2.2.1 Metrics

The metrics used were the same as the ones used in ViSioNS Benchmark, but they were adapted to group results of different MSS separately.

- **Efficiency**: We measure the proportion of targets found for a given number of fixations and report the area under this curve (AUC), noted as AUCperf [9].
- **Scanpath similarity**: Multi-Match (MM) [11] is a pairwise similarity metric that represents saccades as vectors in a two dimensional space. Then, in this space, scanpaths are sequences of vectors. Two such sequences (which can differ in length) are compared on four dimensions: vector shape, vector length (saccadic amplitude), vector position, and vector direction for a multidimensional similarity evaluation. The temporal dimension was excluded as we were not considering fixations’ duration. For each model, its scanpaths were compared to each participant’s scanpaths (after reducing them to match the model’s scanpaths dimensions) and the outcome was averaged across all dimensions and participants, resulting in a single scanpath similarity value for each image (human-model Multi-Match or hmMM). The same was done with the participants’ scanpaths, comparing them within themselves, and thus obtaining a human ground truth for each image (within-human Multi-Match or whMM). These operations were performed for every scanpath with length greater than two in which the target was found. We report AvgMM as the mean value of hmMM, in the case of models, and whMM, in the case of participants, over all images.
- H**uman scanpath prediction (HSP)**: For a given participant’s scanpath, we evaluated how well each model predicted the next human fixation based on the scanpath history [18]. The key idea behind this method is to force the models to follow the participant’s scanpath (ignoring its own predictions). At each step, each model creates what is called a “conditional priority map” (a priority map based on the participant’s previous fixations from where the next fixation is sampled) and we compared the position where the model’s fixation would land (i.e. its prediction of the next fixation) against the participant’s fixation [18]. By using the latter as the ground truth, this allowed for the computation of well-established metrics (such as AUC [7], NSS (Normalized Scanpath Saliency) [28], and Information Gain [19] relative to the center bias and uniform models; we refer to their average across fixations as AUChsp, NSShsp, IGhsp and LLhsp, respectively.

#### 2.2.2 Control conditions

In this project we wanted the model to behave like humans, which was why their cumulative performance and multimatch were the ground truth for our models.

The other three control conditions were the ones used in [36], which are similar to those in [18]. Their goal was to provide a lower and upper bound to the performance of the evaluated models. The lower bound was determined by the uniform and center bias models, and the upper bound by the Gold Standard model (GS). The uniform model predicts fixations to be uniformly and independently distributed over the image. The center bias model stems from [34], who show that people have a tendency to look at the center of images. As this phenomenon occurs mainly during free-viewing experiments [8], we used a Gaussian Kernel Density Estimate (GKDE) over all publicly available fixations in all images of the CAT2000 training dataset [3], excluding the first fixation as it was forced. Lastly, the Gold Standard model predicted fixations of each participant by means of a GKDE over all fixations of other participants on the same image [19]. In these last two cases, the bandwidth of the GKDE has been selected to yield maximum log-likelihood in a 5-fold cross validation paradigm. Since the baseline models are not visual search models themselves (i.e. they don’t generate scanpaths), only the fixation-by-fixation metrics (HSP) were computed on these.

Finally, in order to sort the models by a compound variable, we defined their score as the average of the individual normalized metrics, relative to Humans in the case of efficiency and scanpath similarity metrics (AUCperf, AvgMM) or by the Gold Standard in the case of the scanpath prediction metrics (AUChsp, NSShsp, IGhsp, LLhsp). For AUCperf, this was done by performing *−*|*AUCperf*_*Model*_*−AUCperf*_*Humans*_|, as we intended to maximize similarity to human performance. The rest of the components of the score were computed as (*V alue*_*Model*_ *− V alue*_*Reference*_)*/V alue*_*Reference*_, where Reference stands for Humans in AvgMM and for GS in all other cases. Corr did not have a reference value, so *Corr−* 1 was the component used in the score. Notice that a value of 0 for each and every component of the score means that the model was behaving like humans according to the corresponding metric. A score of 0 means that a model behaved like humans according to every metric.

#### 2.2.3 nnELM model structure

The [6] model consists of a Bayesian Framework that incorporated information iteratively with each saccade (see Fig. 2 for an overview). The posterior was computed as follows:

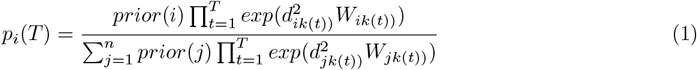

W is what we called “Visual Evidence” (template response in [5]) whose values were sampled from a 2D normal distribution with mean and variance as follows:

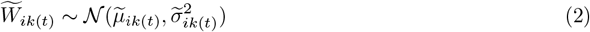

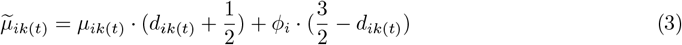

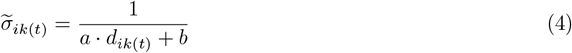

where in both equations k(t) is the fixation at step t; i and j are locations in the image; *ϕ*_*i*_ is a number between -0.5 and 0.5; *µ*_*ik*(*t*)_ is either -0.5 or 0.5; and d is what we called visibility map (also called detectability or eccentricity).

**Figure 2.**
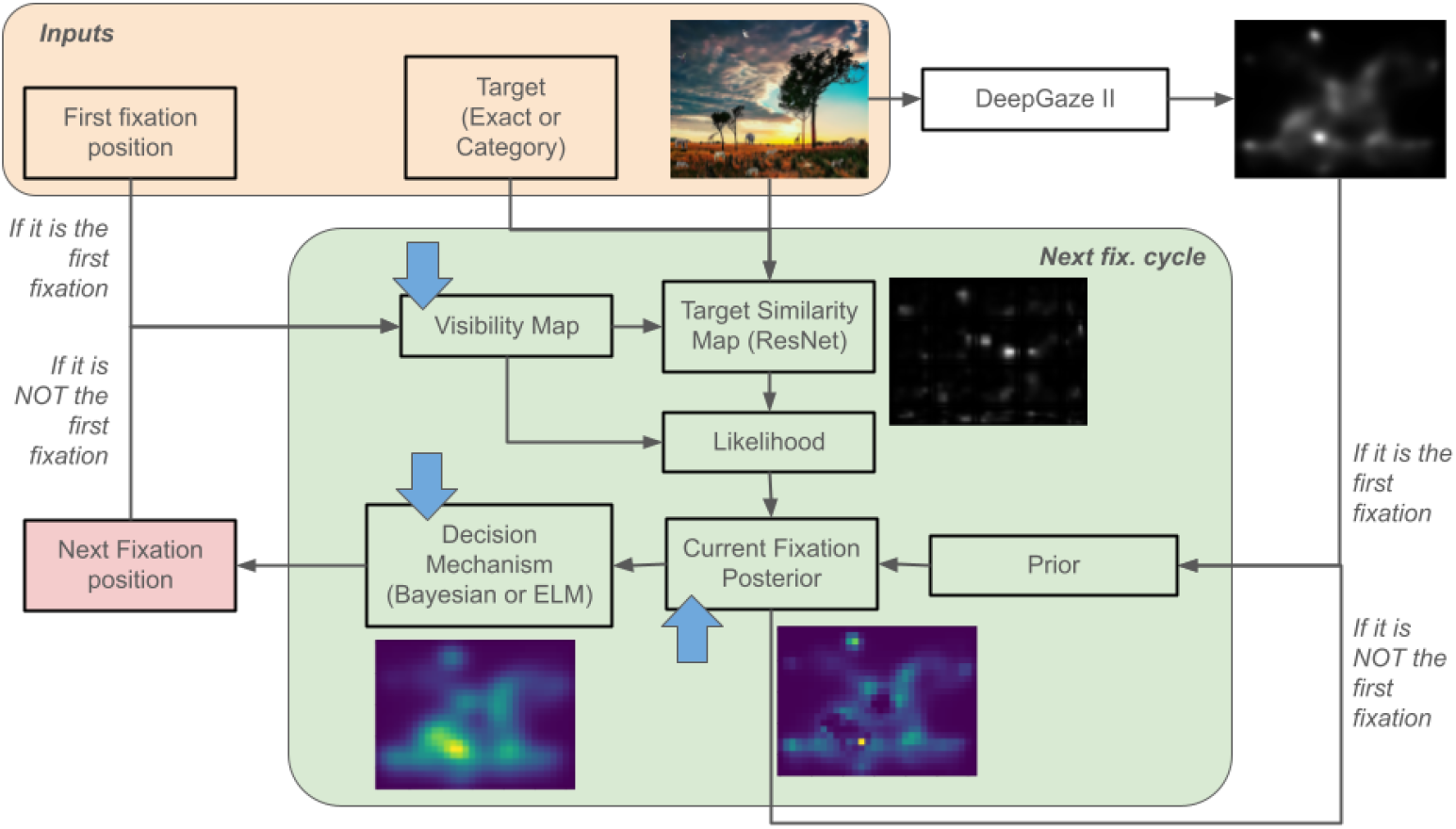
The model receives the image, the target and the position of the first fixation. Before it starts moving it calculates the first prior using the DeepGaze II model which gives the first gist of the image. The model estimates the next fixation position which updates the input values. Also, the posterior becomes the prior of the next cycle. Blue arrows indicate where the model was updated in the current version: A new version of the visibility map is presented, a limited working memory is introduced in the Posterior estimation, and a decision criterion is used to adapt the model to Hybrid Search.

**Figure 3.**
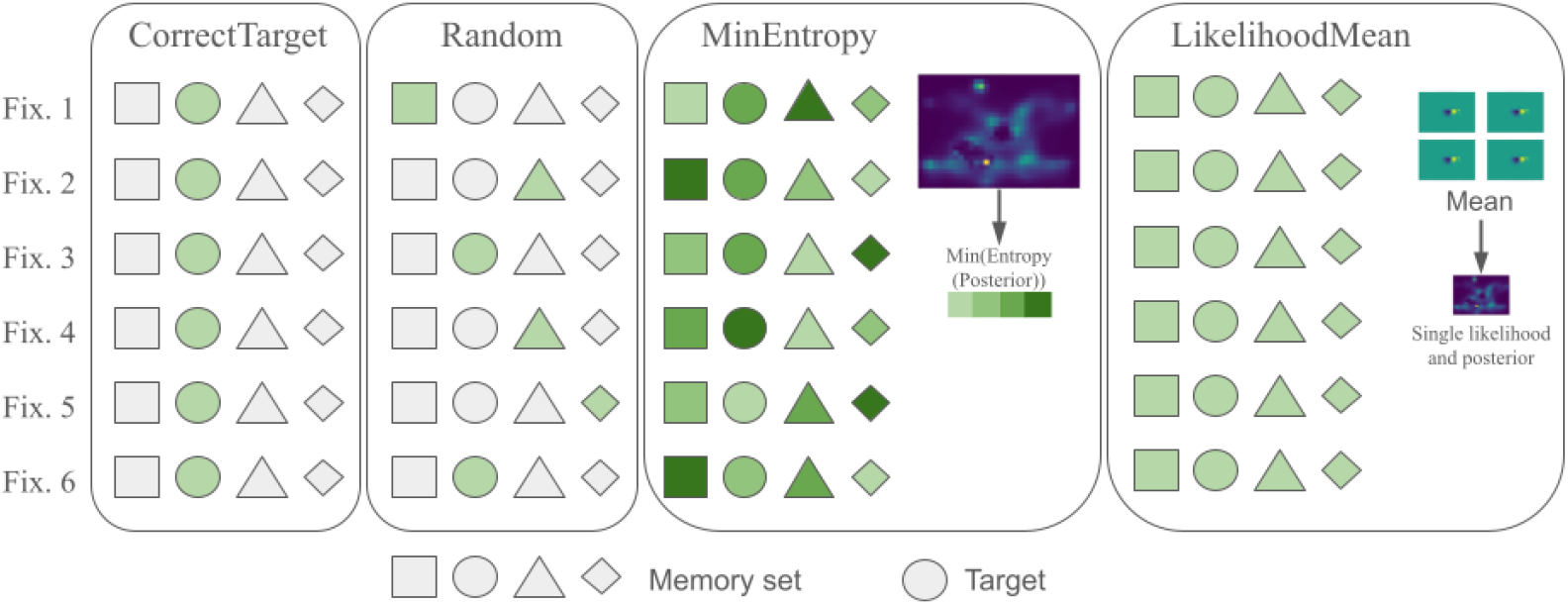
Overview of the selection criteria. CorrectTarget, Random and MinEntropy perform a selection while LikelihoodMean does not.

The prior in the first step of the iteration is a saliency map generated with Deepgaze II from [17]. It is akin to the visual information a human would acquire at first glance. d is the same 2D Gaussian that was used in [5]. The difference in W between this work and the latter is that phi is now computed the same way the attention map is computed in [44], but with ResNeXt-101 32×8d [41] instead of VGG-16 [33].

With the posterior and d, the next fixation is computed the same was as in [6]

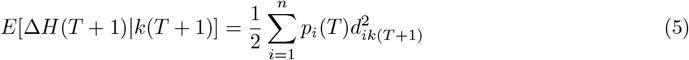

This equation comes from [22]. Theoretically speaking they made assumptions that do not necessarily apply to this setup of natural scenes, but empirically speaking it significantly reduced processing times without compromising search efficiency, enhancing the model’s real-time utility and hastening all subsequent experiments and analyses.

### 2.3 Motivation for changes and explanation

#### 2.3.1 Peripheral vision

In order for the likelihood to carry information from the peripheral vision while keeping the IOR restricted to the foveal region we replaced the term 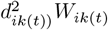 in Eq.1 by:

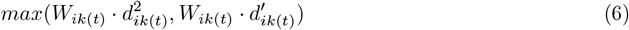

where:

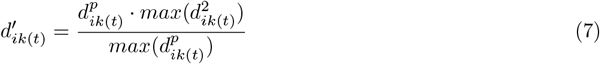

and 0 *< p*≤ 1 was a parameter to explore. *W*_*ik*(*t*)_ *d*_*ik*(*t*)_ kept the same values as before: if the target was not within the fovea, the values in the foveal region would be negative and it would have positive values otherwise. Outside of the fovea it could have values of 0 or near 0. Meanwhile 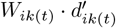 had the same principle but with more variance.

If we took values of both terms for any given location and they were both negative, nothing would happen because the values in 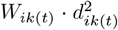 would always be closer to 0 than the 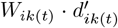 values.

Thus 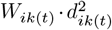 would be chosen by the max operation. In contrast, if both values were positive (only possible within the fovea), 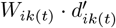 have a greater value than 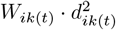 , so 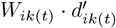 would survive the max operation. This could only happen if the target was in the current location.

It could not happen that either term was positive while the other was negative. It could not happen that 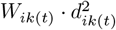 was non-zero while 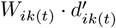 was zero. The opposite could have happened and in that case, 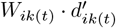 would survive the max operation if it was positive, and 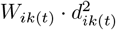 otherwise. Both of these scenarios could only take place in the peripheral area. This means that if an object similar to the target object (a distractor) was in the peripheral area, it would be taken into account by the model.

All in all, this modification provided an attention mechanism for the whole field of vision while keeping the inhibition of return applied to the fovea only. 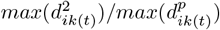 was the term that allowed all of these behaviours at the same time.

The term *d*^2^ was also used in eqs. 3, 4 and 5. As for eqs 3 and 4, *W*_*ik*(*t*)_ was sampled from a normal distribution and both *µ*^*′*^ and *σ*^*′*^ depend on *d. ϕ* in eq. 3 is the target similarity map: it captures how similar the target is to each region of the image according to a neural network. *µ* is a mask defined by:

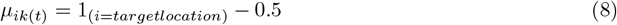

so it knows exactly where the target is. The formula in eq. 3 gives a lot of weight to the real target within the fovea and less weight elsewhere (first addend). At the same time it gives more weight to the real target and distractors in the peripheral vision and less weight in the fovea (second addend). That is to say, it will only pay attention to an object within the fovea if it is indeed the real target, but it can also pay attention to other objects that are not within the fovea.

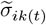 indicates how much noise there is: the idea is that we want little noise within the fovea and more noise as we move further away.

Previously these effects in mu and sigma did not matter much in the sense that everything in the peripheral vision was discarded afterwards, so only the desired effects within the fovea were what mattered. After the change in eq. 1 the information from the peripheral vision started to be considered as well, so both of these formulas started mattering much more. Still, there was a distinction: in 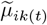 we wanted to distinguish distractors from the real target, but by definition, they are called distractors because we are not sure whether they are the real target or not until we look at them, so we only cared about making a clear distinction between target and distractor within the fovea. Then it made sense to Use 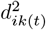 in eq. 3 instead of 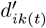 :

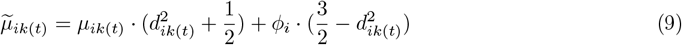

On the other hand, we cared about 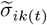 all over the visual field, so it made sense to use 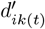 in eq.

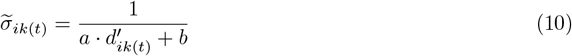

In Eq. 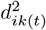 we did not want a bigger foveal region so no changes were made.

#### 2.3.2 Visual working memory

We did not include the modifications to the peripheral vision in the following equations to maintain simplicity.

In visual search and hybrid search, the visual working memory is the amount of fixations being remembered while performing the search. The first strategy to limit the model’s visual working memory was to take the last N fixations and completely forget the rest, like what was done in [45]. Eq 1 changed as follows:

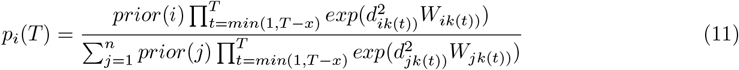

where x is the amount of fixations that the model remembers.

This brought the following variant into the fold: if the model forgets fixations it made, it might made sense to forget the initial prior as well. The equation remained the same when *T≤ x*, but when *T > x* the equation changed as follows:

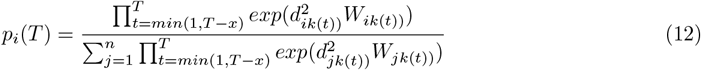

We also developed another approach in which instead of completely forgetting fixations we applied an exponential decay to the working memory. The idea was to progressively remember fixations less and less as the search went on. The parameters of the exponential were selected so that after approximately 8 fixations, the model will almost completely forget about the current one:

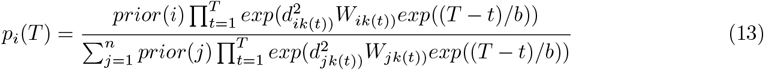

this version could also forget the prior like this:

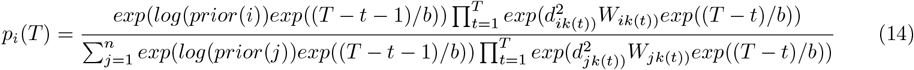

where in both cases *b >* 0 was a parameter to explore.

#### 2.3.3. Hybrid task adaptation

In order to make IBS work in hybrid tasks, we had to allow it to process more than one object. We did that by computing the likelihood and posterior corresponding to each object in the memory set at each step. Those posteriors would have to be somehow combined or selected. To do so, we implemented four strategies so far:

- Random: Selecting the posterior of a random object.
- MinEntropy: Selecting the posterior with less entropy (it can be a 2D or regular entropy).
- CorrectTarget: Selecting the posterior of the real target. This one served as a baseline.
- LikelihoodMean: Taking the mean of the foveated visual evidences 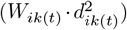 over the memory set and computing a single likelihood and posterior with that.

### 2.4 Validation set and Benchmark

Participants from the Hybrid Search dataset were split up into train and validation sets using the train test split function from Scikit learn Python package^2^ with 21% validation set size and stratified by subject accuracy. This procedure resulted in 34 and 10 participants in train and validation sets respectively.

Also, an external validation was performed using datasets and measures available in the ViSioNS benchmark of visual search in natural scenes ^3^ [36]. Briefly, the Interiors dataset from our previous work comprises 134 images and 57 participants who had to look for an exact target in photos of interiors [5]. The MCS and COCOSearch18 datasets used by Zelinsky and collaborators comprises 1687 and 612 images and 23 and 10 participants respectively [9] that had to look for any target belonging to a given category (for instance, a “cup”) in a varied set of images. The Unrestricted dataset collected by Zhang, Kreiman, and collaborators comprises 234 images and 15 participants who also had to look for a target within a category as MCS and COCOSearch18 [44].

### 2.5 Results

#### 2.5.1 Human behavior

The response time of the participants performing the Hybrid Search followed a logarithmic dependence with the memory set size (MSS), consistent with previous findings [1] (Fig. 4 D). Target detection accuracy decayed with MSS = 4 (Fig. 4 E).

In order to complete the task, participants performed a total of 9.03±0.15, 10.45±0.17, and 12.79±0.19 fixations across different images, for MSS equal to 1, 2, and 4 respectively (Fig. 4 G). Most fixations were on new items, as the mean number of re-fixations were 0.59 ± 0.02, 0.72 ± 0.03, and 1.02 ± 0.03 respectively. The saccade direction distribution presented a bias towards horizontal as is typically seen in natural scene exploration and search (Fig. 4H). The main sequence of saccades (Fig. 4I) was also similar to previous reports in almost every task [25].

**Figure 4.**
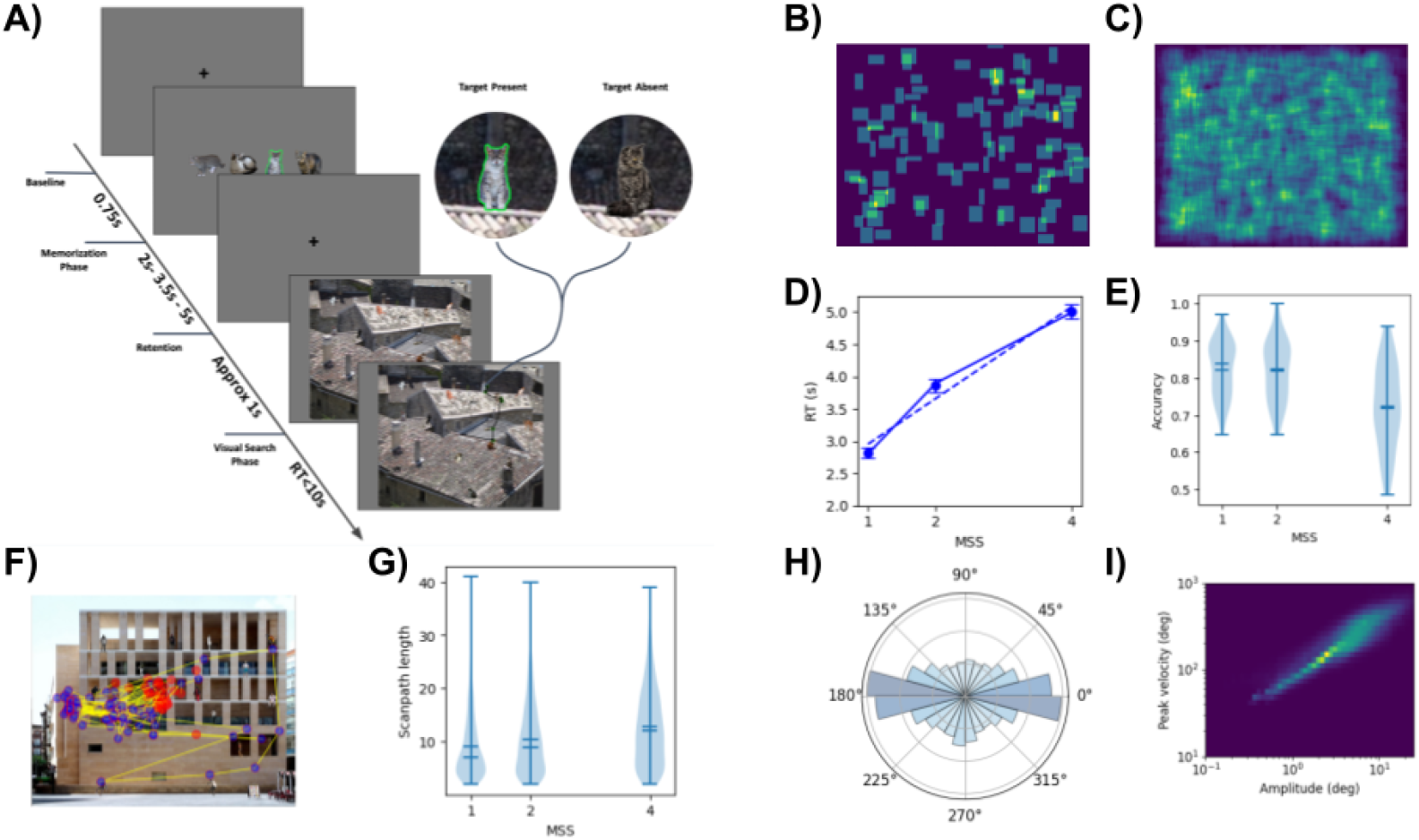
Experimental paradigm. **A**. Time progression of the trial. A detail of a highlighted version of the target at present condition and the distractor item at absent condition. After a fixation dot a variable number of potential targets were presented (Memory Set Size, MSS = 1, 2, 4), followed by a retention period. Search screen always have 16 items (Visual Set Size, VSS), and participant’s task was to respond whether one or none of the potential targets was present. In target absent trial a cat different from the targets is included in the same location. See [1]. **B, C**. Heatmap of the bounding boxes from the items presented at visual search phase across trials. B. Targets, C. Distractors. **D, E**. Performance measured as Response Times (on Target Present trials) and Accuracy (on Target Present trials) as function of Memory Set Size (MSS). **F**. Scanpaths of all participants in a sample image. **G**. Scanpath length as function of MSS. **H**. Saccade direction. **I**. Main sequence of saccades.

#### 2.5.2 Improving visual search model’s performance: Effects of Visibility and Working memory capacity

Before extending the model from Visual Search (MSS=1) to Hybrid Search (for trials with MSS = 2 and 4), we focused on improving the Visual Search model. Recently, [6] proposed using the Entropy-Limit Minimization (ELM) as a model of decision and guidance (see also [45, 29]). We suggested two modifications to achieve a greater similarity between the model and humans: Weighting the tails of the visibility map distribution to model the peripheral vision and limiting the visual working memory capacity. The previously defined train set (79%) was used to tune the parameters of the modifications and both the validation set (21%) and the ViSioNS benchmark [36] were used to validate those parameters. Until [6] the model acquired new information from the fovea only (see Fig. 5). Since the visibility map followed a Gaussian distribution, the likelihood value in the peripheral region remained constant at 1 (given that *exp*(0) = 1), which was non-informative. In contrast, the foveal region could display positive values. Values larger than 1 in a region within the fovea suggested the presence of a relevant feature, and most likely the model would have fixated on that location next, especially if it had been close to the target already. When values were close to 0, it indicated that there was no relevant information, which elicited an inhibition of return across the entire fovea. After this happened, the model would rely on the prior to guide the next fixation. Recursively, if an inhibition of return had been applied to every fixation, then the saliency map (initial prior) would have dictated the location of subsequent fixations. To improve this approach, we made the likelihood more informative by allowing it to capture information from the peripheral vision. We modified eq. 1 by dividing the visual evidence into foveal and peripheral terms that compete with each other according to a max operation, which also kept the inhibition of return restricted to the foveal region (see eq. 6). This led to the likelihood displaying values larger than 1 (see Fig 5 A). We also replaced the term *d* by *d*^2^ in eq. 3, and *d* by *d*^*′*^ in eq. 4 (see eqs. 9, 10). These changes made the model more consistent with what is currently known about the foveal and peripheral areas.

**Figure 5.**
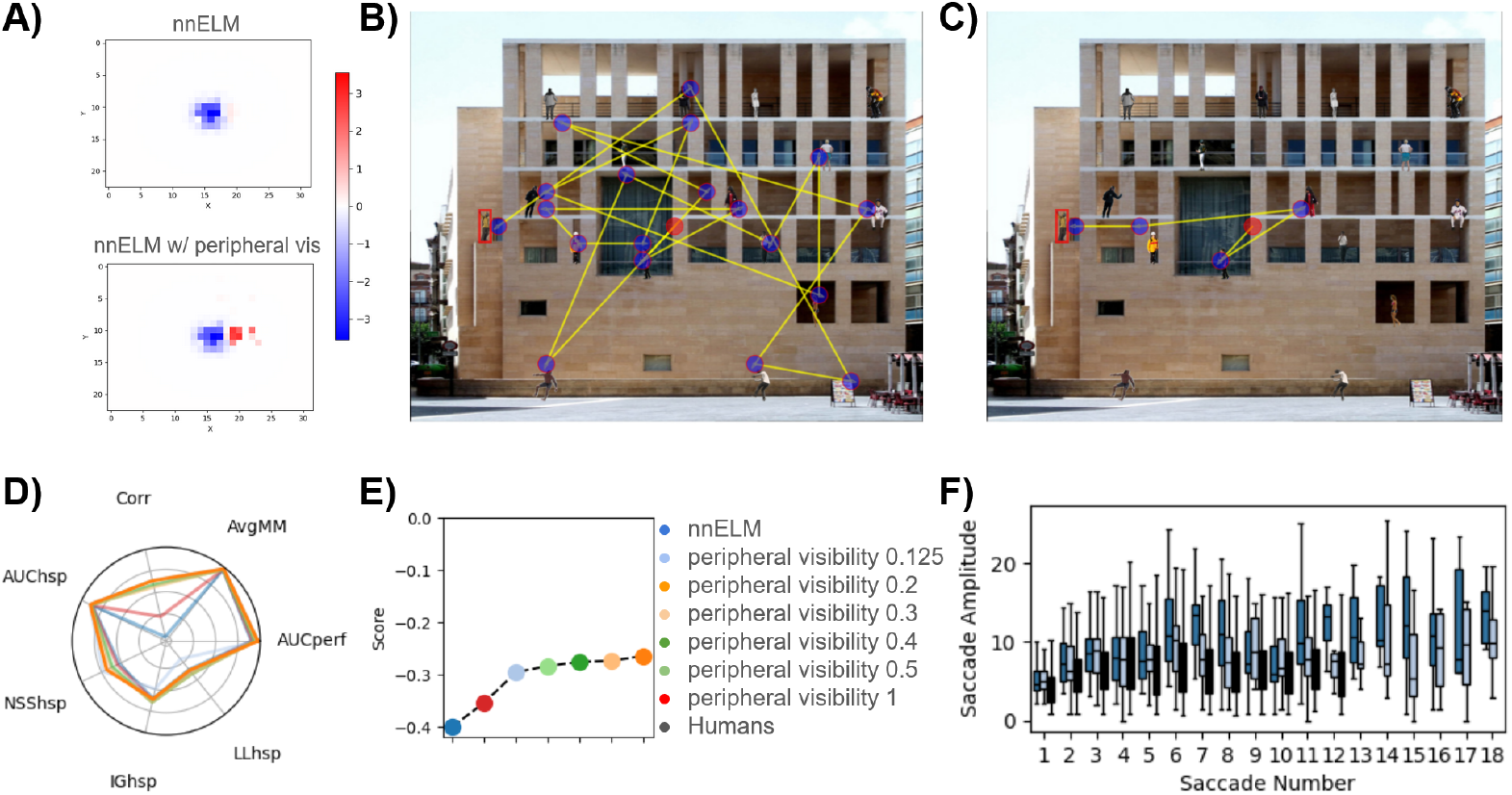
Peripheral Visibility. **A**. Change in likelihood after adding peripheral visibility to the model. **B**. Model scanpath before adding peripheral visibility to the model. **C**. Model scanpath after adding peripheral visibility to the model. **B**,**C**. The red dot is the initial fixation, the blue dots are the other fixations and the yellow lines are saccades. **D**. Model performance according to each metric. With p = 0.2 the model has a higher value for almost every metric. **E**. Model score. With p = 0.2 the model has a greater score than the rest. **F**. Saccade amplitude boxplots for the model with and without peripheral visibility, and how they compare against humans, only for trials in which the target was found. 90% quantile of the saccade number was used to filter longer scanpaths.

This model outperformed the nnELM model [6] in every metric (Fig 5 D). In order to choose a value for the exponent (p in eq. 7) we found a compromise between more general metrics like the AUCperf and MM and the fixation-based metrics, in particular IG and LL. Thus, the best model had *p* = 0.2 (Fig 5 E). It is important to note that small changes in the exponent did not significantly affect the results. In a previous version of the model, we observed that after some time the saccades tended to be longer, presumably because the saliency map was what primarily dictated where the model went. By improving the visibility map the saccade amplitude also got closer to human saccades along the whole scanpath (Fig. 5 B, C, F), without any ad-hoc constraint on the saccade amplitude [45].

Human participants have memory limitations that makes them less likely to remember which regions have already been visited in long searches. Meanwhile, previous IBS and ELM models [5, 36, 6] had unlimited memory capacity and would eventually find the target if the scanpath was large enough. To address this discrepancy, we implemented memory constraints within the model, thereby bringing its behavior into closer conformity with that of human participants during extended searches. Because of that, we decided to take into account only the last N fixations and completely forget the rest, like what was done in [45] (see eq. 10).

We had a non-trivial prior, which is why it was important to decide whether the model could forget the prior or not as if it was another fixation. Firstly, we took the best model from the previous experiment, made it forget the prior and explored the number of fixations that it could maintain in visual working memory (see eq. 11). In this case all limited memory models outperformed the unlimited memory model, with 4 being the number of fixations that worked best (Fig. 6 A, B). It is worth mentioning that the metrics’ values were very close to those of the models with a visual working memory of 6, 8 and 12 fixations (Fig. 6 A, B), which is consistent with [45], who find good values to be between 4 and 12 fixations.

**Figure 6.**
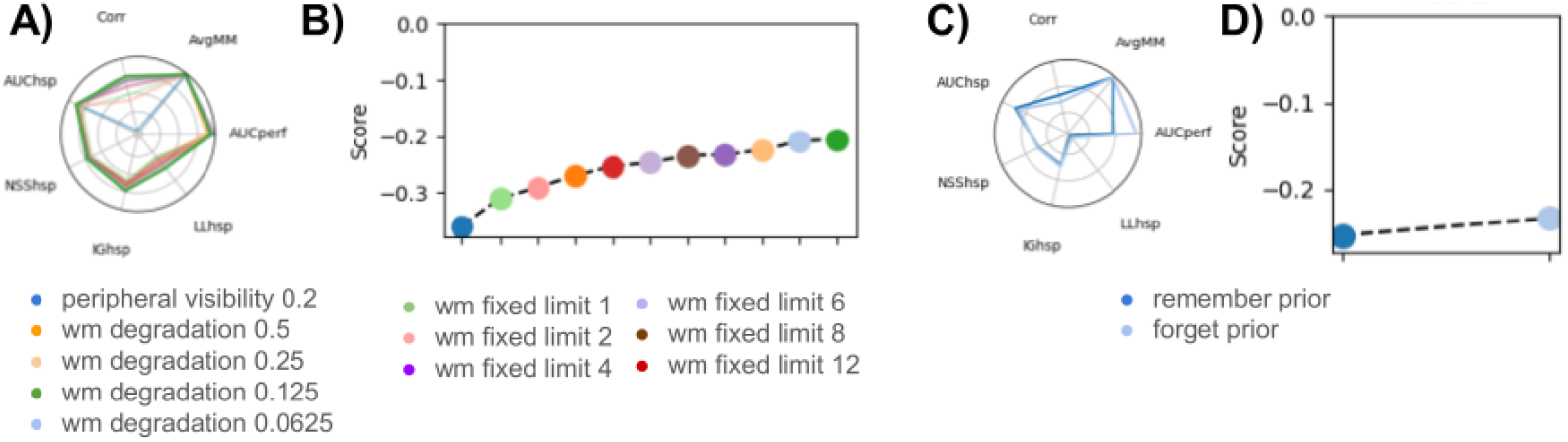
Limited Memory. **A**. Model performance according to each metric. Degrading the memory with an exponential (exponent = 0.125) works best for almost every metric. **B**. Model score. An exponent of 0.125 yields the highest score. **C**. Forgetting the prior doesn’t yield better results with the exception of corr and AUChsp. **D**. The score remains higher when forgetting the prior.

Secondly, models of other memory domains suggest a decay in the retention of previous items instead of a fixed all-none limit [35]. Thus, we replaced the stepwise function by an exponential decay and explore the time constant. We found that the best model has a decay of 0.125, which corresponds to a decrease of 0.4 for 4 fixations and 0.63 after 8 fixations (Fig 6 A, B). This worked even better than the fixed-limit models.

Finally, we took the best model and made it remember the prior, and found that it was worse overall (Fig. 6 C, D).

#### 2.5.3 Adapting the model to Hybrid search: Target selection criteria

For the rest of the experimentation we proceeded with the best model identified so far, which included an improved visibility map (with an exponent of 0.2) and an exponential decay of previous fixations (with an exponent of 0.125 and considering the prior as fixation zero). For a single target, the model determined where to look next based on a single probability map. However, when dealing with many potential targets, the observer has to decide at each step which map to use or how to combine them. Here we propose that the posterior probability map for every target is estimated in parallel, and the model guides the next fixation towards the map that contains the largest amount of information (i.e., minimum entropy). We compared this criterion with different alternatives, such as selecting maps at random in each fixation or knowing the correct target in advance and only focusing on that map (see Fig. 7 A). The MinEntropy approach worked better for memory set sizes of 2 and 4 (see Fig 7 B, C).

**Figure 7.**
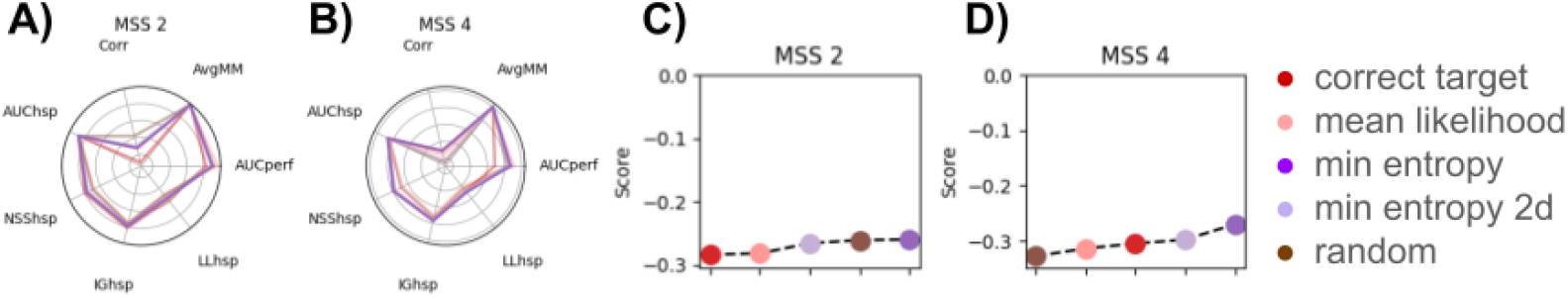
Hybrid Search. **A, B**. Model performance according to each metric for MSS 2 trials and MSS 4 trials respectively. MinEntropy works better for almost every metric. **C, D**. Model score for MSS 2 trials and MSS 4 trials respectively. MinEntropy yields the highest score in both cases.

#### 2.5.4 Model validation

A validation set stratified by accuracy was separated at the beginning of the experiment. The best model also showed a better performance with the test participants for the three MSS values (Fig 8).

**Figure 8.**
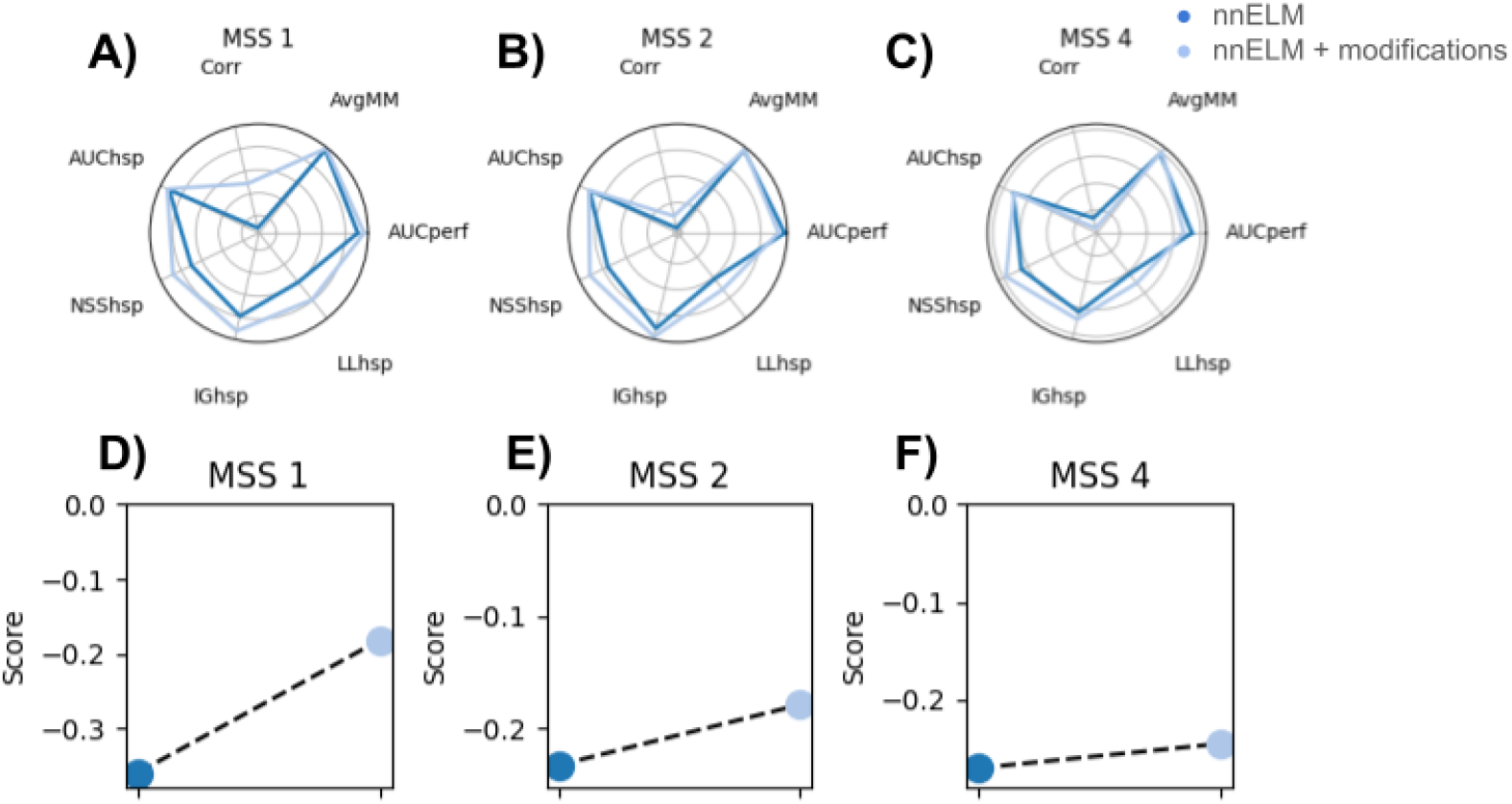
Validation. **A, B, C**. Model performance according to each metric for MSS 1, 2 and 4 trials respectively. The modifications improve every metric for MSS 1, and almost every metric for MSS 2 and 4. **D, E, F**. model score for MSS 1, 2 and 4 trials respectively. The modifications improve the overall score for every MSS.

A validation set stratified by accuracy was separated at the beginning of the experiment. The best model also showed a better performance with the test participants for the three MSS values (Fig 8A). In the case of nnELM we used the CorrectTarget strategy for MSS 2 and 4 because originally it could not perform hybrid search tasks. Moreover, our best model showed better performance in every metric than previously published versions such as nnIBS and other state-of-the-art models. Table 1 corresponds to the average across the four datasets of the ViSioNS Benchmark (to date these are: Unrestricted, Interiors, COCOSearch18 and People) [36]. This result lowered the possibility of overfitting to the HSEM dataset.

**Table 1:** Benchmark evaluation. Average across all datasets.

| Model | AUCperf | AvgMM | Corr | AUCshp | NSShsp | IGhsp | LLhsp | Score |
| --- | --- | --- | --- | --- | --- | --- | --- | --- |
| Human performance | 0.617 | 0.871 | - | - | - | - | - | 0.0 |
| gold standard baseline | - | - | - | 0.908 | 2.922 | 2.746 | 2.248 | 0.0 |
| nnELM + mods | 0.673 | 0.852 | 0.256 | 0.807 | 1.801 | 1.872 | 1.115 | -0.303 |
| nnELM | 0.52 | 0.843 | 0.219 | 0.796 | 1.445 | 1.623 | 0.902 | -0.364 |
| nnIBS | 0.507 | 0.843 | 0.228 | 0.799 | 1.444 | 1.22 | 0.499 | -0.411 |
| Gupta, et al. (2021) | 0.465 | 0.828 | 0.169 | 0.681 | 1.006 | -0.866 | -1.312 | -0.691 |
| uniform baseline | - | - | - | 0.5 | 0.0 | 0.499 | 0.0 | -0.817 |
| center bias baseline | - | - | - | 0.66 | 0.543 | 0.0 | -0.499 | -0.827 |
| Zhang, et al. (2018) | 0.569 | 0.79 | 0.122 | 0.619 | 0.85 | -2.054 | -2.528 | -0.846 |
| Yang, et al. (2020) | 0.3 | 0.599 | 0.058 | 0.49 | 0.897 | -3.053 | -3.366 | -1.048 |

**Table 2:** Ranking MSS 1.

| Model | AUCperf | AvgMM | Corr | AUCshp | NSSshp | IGhsp | LLhsp | Score |
| --- | --- | --- | --- | --- | --- | --- | --- | --- |
| Human performance | 0.598 | 0.841 | - | 0.861 | 2.129 | 2.078 | 1.495 | 0.0 |
| gold standard baseline | - | - | - | - | - | - | - | - |
| wm degrad 0.125 | 0.654 | 0.831 | 0.751 | 0.761 | 1.539 | 1.538 | 0.808 | -0.204 |
| wm degrad 0.0625 | 0.606 | 0.826 | 0.714 | 0.765 | 1.516 | 1.51 | 0.779 | -0.209 |
| wm degrad 0.25 | 0.699 | 0.834 | 0.753 | 0.75 | 1.529 | 1.467 | 0.736 | -0.224 |
| wm fixed limit 4 | 0.635 | 0.834 | 0.68 | 0.761 | 1.478 | 1.449 | 0.718 | -0.232 |
| wm fixed limit 8 | 0.625 | 0.826 | 0.697 | 0.759 | 1.473 | 1.417 | 0.686 | -0.233 |
| wm fixed limit 6 | 0.656 | 0.829 | 0.614 | 0.753 | 1.474 | 1.439 | 0.709 | -0.246 |
| forget prior | 0.377 | 0.834 | 0.739 | 0.764 | 1.462 | 1.424 | 0.693 | -0.252 |
| wm fixed limit 12 | 0.598 | 0.824 | 0.612 | 0.763 | 1.464 | 1.351 | 0.662 | -0.253 |
| per vis 0.200 | 0.573 | 0.826 | 0.639 | 0.767 | 1.462 | 1.273 | 0.542 | -0.264 |
| wm degrad 0.5 | 0.712 | 0.837 | 0.7 | 0.729 | 1.439 | 1.3 | 0.569 | -0.27 |
| per vis 0.300 | 0.558 | 0.821 | 0.565 | 0.761 | 1.377 | 1.343 | 0.612 | -0.273 |
| per vis 0.400 | 0.542 | 0.817 | 0.593 | 0.757 | 1.324 | 1.348 | 0.618 | -0.275 |
| per vis 0.500 | 0.538 | 0.814 | 0.56 | 0.754 | 1.285 | 1.34 | 0.609 | -0.283 |
| wm fixed limit 2 | 0.673 | 0.832 | 0.416 | 0.716 | 1.467 | 1.396 | 0.665 | -0.29 |
| per vis 0.125 | 0.619 | 0.829 | 0.643 | 0.772 | 1.559 | 1.031 | 0.3 | -0.299 |
| wm fixed limit 1 | 0.625 | 0.831 | 0.501 | 0.685 | 1.376 | 1.235 | 0.504 | -0.309 |
| per vis 1.000 | 0.51 | 0.816 | 0.226 | 0.747 | 1.181 | 1.293 | 0.506 | -0.353 |
| nnELM | 0.573 | 0.814 | -0.008 | 0.767 | 1.462 | 1.273 | 0.542 | -0.359 |
| uniform baseline | - | - | - | 0.5 | 0.0 | 0.583 | 0.0 | -0.785 |
| center bias baseline | - | - | - | -0.646 | 0.562 | 0.0 | -0.583 | -0.844 |

**Table 3:** Ranking MSS 2.

| Model | AUCperf | AvgMM | Corr | AUCshp | NSSshp | IGhsp | LLhsp | Score |
| --- | --- | --- | --- | --- | --- | --- | --- | --- |
| Human performance | 0.548 | 0.838 | - | - | - | - | - | 0.0 |
| gold standard baseline | - | - | - | 0.855 | 1.983 | 1.875 | 1.384 | 0.0 |
| min entropy | 0.645 | 0.829 | 0.283 | 0.756 | 1.543 | 1.487 | 0.76 | -0.26 |
| random | 0.633 | 0.83 | 0.427 | 0.748 | 1.392 | 1.426 | 0.699 | -0.261 |
| min entropy | 0.657 | 0.83 | 0.313 | 0.75 | 1.49 | 1.462 | 0.736 | -0.266 |
| likelihood mean | 0.51 | 0.829 | 0.269 | 0.748 | 1.357 | 1.41 | 0.683 | -0.282 |
| correct target | 0.722 | 0.834 | 0.107 | 0.758 | 1.593 | 1.527 | 0.8 | -0.284 |
| nnELM | 0.561 | 0.816 | 0.131 | 0.752 | 1.235 | 1.351 | 0.624 | -0.319 |
| uniform baseline | - | - | - | 0.5 | 0.0 | 0.491 | 0.0 | -0.788 |
| center bias baseline | - | - | - | -0.656 | 0.658 | 0.0 | -0.491 | -0.814 |

**Table 4:** Ranking MSS 4.

| Model | AUCperf | AvgMM | Corr | AUCshp | NSSshp | IGhsp | LLhsp | Score |
| --- | --- | --- | --- | --- | --- | --- | --- | --- |
| Human performance | 0.446 | 0.842 | - | - | - | - | - | 0.0 |
| gold standard baseline | - | - | - | 0.854 | 1.813 | 1.859 | 1.325 | 0.0 |
| min entropy | 0.573 | 0.843 | 0.297 | 0.741 | 1.437 | 1.436 | 0.668 | -0.27 |
| min entropy 2d | 0.595 | 0.842 | 0.181 | 0.732 | 1.395 | 1.419 | 0.651 | -0.298 |
| correct target | 0.757 | 0.841 | 0.291 | 0.735 | 1.415 | 1.414 | 0.646 | -0.305 |
| nnELM | 0.5 | 0.818 | 0.193 | 0.744 | 1.234 | 1.331 | 0.563 | -0.314 |
| likelihood mean | 0.534 | 0.831 | 0.244 | 0.721 | 1.219 | 1.331 | 0.563 | -0.314 |
| random | 0.606 | 0.835 | 0.153 | 0.725 | 1.258 | 1.361 | 0.593 | -0.328 |
| uniform baseline | - | - | - | 0.5 | 0.0 | 0.534 | 0.0 | -0.782 |
| center bias baseline | - | - | - | -0.654 | 0.587 | 0.0 | -0.534 | -0.828 |

## 3 Discussion

This work introduces a novel computational model for Hybrid Search tasks in natural scenes, extending the capabilities of existing models focused solely on Visual Search. By incorporating mechanisms like limited memory capacity and an adapted visibility function, our model effectively captures aspects of human behaviour observed in hybrid search experiments. These adaptations allowed the model to perform better than previous approaches within the visual search benchmark [36]. This work conveys the release of a new Hybrid Search Eye Movements (HSEM) dataset, with 4,442 scanpaths from 44 participants performing a hybrid visual and memory search where targets and distractors were superimposed to natural scenes.

Our model’s performance demonstrates the importance of peripheral visibility and visual working memory (VWM) in hybrid search tasks. Previous studies have noted that human searchers utilise peripheral information even as the foveal region remains the primary source of high-resolution detail [2]. In particular, the periphery has an important role in the guidance of eye movements by incorporating information of the context [REFS]. For instance, [27] showed that scene context acts as a framework to guide our vision, particularly in peripheral vision. Nuthmann and Malcom [24] highlighted the importance of

colour in the periphery to help localize targets. By enhancing our visibility map to reflect peripheral information, we achieved significant improvements in alignment with human-like saccadic behaviour and fixation patterns, particularly in scenarios with higher memory set sizes. Geisler and collaborators also stressed the importance of peripheral vision in search guidance [21, 13, 4]. In their paradigms, they made a strong effort to estimate the visibility map for each participant. They used the first 16 sessions only to measure the dependence of the visibility with the eccentricity in each direction. Those measurements were then used to estimate the parameters of the IBS’s visibility map for each participant [21]. This approach was also used in the recent paper by [45] for the same kind of artificial stimuli. While this approach could potentially incorporate individual differences among participants, their generalization capability might be more limited. In contrast, our approach avoids a potential leak of information about the viewing patterns from the participants to the model because its parameters are not fitted to each participant [5, 15]. Future works could systematically measure the distribution of eccentricities from different participants across different stimuli, including natural scenes, and use those distributions to train a DNN model capable of producing comparable visibility maps.

Visual Working Memory (VWM) is a core mechanism in our visual system. We evaluated two approaches to limit the VWM capacity: adding a fixed amount of fixations or an exponential decay to the model’s memory. Zhou and Yu [45] implemented a model for visual search with limited VWM in artificial displays. Interestingly, they found the optimal value to be around 8 fixations, coinciding precisely with our modeling results. Moreover, this goes in line with behavioural results of Kaunitz and collaborators [16], who found that participants remembered up to seven fixated non-target faces with more than 70% accuracy in an incidental memories paradigm. Our second approach, incorporating an exponential decay to the model’s memory, improved the model’s performance even further, suggesting that the memory of recently explored locations exponentially decays over fixations. This resembles other memory systems, such as iconic memory, in which the exponential decay seems to be the best model [35]. Importantly, comparisons with other systems are relevant as the hybrid search task we studied differs from a standard working memory paradigm, in which the stimulus disappears after a short presentation. In hybrid search, the stimuli and their context remain on the screen and it is the active vision process that generates a sequence of foveated images.

In this work, we generated the similarity map with a ResNeXt-101 32×8d network [41] instead of the VGG-16 network [33] used in previous works [44, 36]. This novel neural network shows better alignment with visual cortex activity (BrainScore^4^), and it can potentially extract more informative features from the images, which might have been missed by earlier architectures. While this network yielded an increase in some of the metrics, their effect sizes were low. A key aspect that these approaches share is that they take into account the visual cues that come from the scene context. However, they fall short in accounting for scene semantics or, in other words, the spatial relationship between objects [30]. For example, if we are looking for a cup, we would typically not look on the floor but instead focus on likely locations such as a table or the countertop. This spatial relationship can be more complex: people are often near cars at ground level, but that is not the case if the person is on a balcony. This information has been slowly learnt through experience. Therefore, adding contextual knowledge that captures scene semantics represents a natural direction for advancing the model’s capabilities.

A critical aspect of hybrid search relies on the decision of how to prioritize possible targets in each fixation. This prioritization could be influenced by both the current state of the search and biases generated during the memorization phase. Previous studies have modelled this process as competing drift-diffusion models, in which it is possible to introduce modulations in both the evidence accumulating rate and the initial bias [12]. However, these models did not take into account the sequential nature of the search, where each fixation represents a decision point in which accumulated evidence may be re-evaluated. Here, we modelled how this decision is taken at each step. The logarithmic dependence of reaction time on memory set size suggests that memory search operates in parallel [12]. Building on this, we assumed that the posterior distribution for each target is processed in parallel, followed by a selection process that determines which map will guide the following saccade. We found that a selection process based on the potential information gained after the next saccade outperformed strategies guided either randomly or by a model that is only aware of the real target. This result is in line with evidence-based integration processes proposed by drift-diffusion models of Hybrid Search [12, 1]. Importantly, the model incorporates the discrete nature of cognitive processes [REF Wyble, et al, 2009; Wyble, et al, 2011; vanRullen, 2016 ], where saccades are considered the natural boundaries of attentional episodes [REF Kamienkowski, et al., 2012; Navajas, et al., 2014].

Regarding the biases generated during the memorization phase, one limitation of our approach is that our models assumed all targets were remembered equally. This simplification prevented us from incorporating the increased difficulty that humans experience as memory set size grows, where some targets may be encoded with different strengths during the memorization phase. For instance, primacy and recency effects, typically observed during the encoding of item sequences [14], were not evident in the current paradigm, likely due to the low memory set sizes used. Increasing set value sizes could show some of these effects [1, 38] which could be readily incorporated into future models. Moreover, in this study all potential targets were similar and belonged to the same category, preventing an evaluation of biases arising from differences in target attributes, such as saliency or target-context congruence. New experiments could specifically evaluate how potential targets’ attributes interact during memory encoding, which could provide the basis for extending our model to a more realistic memory encoding framework.

This study unveiled the first computational model for eye movements during hybrid search in natural scenes, providing a robust framework with key components relevant to explaining human behaviour in these tasks. Additionally, we presented the Hybrid Search Eye Movements (HSEM) dataset, which includes a large variety of images under different conditions, and the corresponding human scanpaths. Using the ViSioNS Benchmark, we validated the enhancements done to our model, demonstrating that our new version outperformed existing models [44, 42, 15]. Overall, this work illustrates how cognitively inspired model adjustments can lead to a more accurate representation of human search behaviour across complex tasks.

## 4 Acknowledgements

J.E.K received research grants from CONICET (PIP 11220150100787CO) and ANPCyT (PICT 2018-2699). J.E.K. and M.J.I. received an award from ARL (Cooperative Agreement Numbers W911NF1920240 and W911NF2120237). We thank Alessandra Barbosa for their collaboration with the data acquisition, and Joaquin Gonzalez and Anthony Ries for insightful discussions.

## 5 Author contributions

GR implemented the computational models. GB developed the ELM version of the IBS model and contributed to the construction of the experimental stimuli. DC preprocessed the eye-movement data and contributed to the construction of the experimental stimuli. MJI and JEK conceptualised the study, designed the experiment, and supervised the project. GR, MJI and JEK analyzed the data and wrote the manuscript.

## 7 Code and data availability

Upon acceptance, the code will be at https://github.com/NeuroLIAA/ruarte-hs-2024 and the data will be shared through OSF.

## Footnotes

1 https://psychopy.org/api/iohub/index.html

2 https://scikit-learn.org/stable/modules/generated/sklearn.model_selection.train_test_split.html

3 https://github.com/NeuroLIAA/visions

4 https://www.brain-score.org/vision/

